# FT-IR micro-spectroscopy for imaging the extracellular matrix composition in biofilms

**DOI:** 10.1101/2024.08.26.609418

**Authors:** Stefan de Bruin, Carina Hof, Mark C. M. van Loosdrecht, Diana Z. Sousa, Yuemei Lin

## Abstract

Microorganisms form granules by embedding themselves in an extracellular matrix through the secretion of extracellular polymeric substances (EPS). The extracellular matrix is a complex structure comprising of e.g. proteins, polysaccharides, lipids, and extracellular DNA. Understanding the function of individual EPS components within the matrix not only requires knowledge on the composition of the extracellular matrix, but also on the spatial distribution of said components. Molecular imaging like e.g. fluorescence microscopy have been used for the visualization of the extracellular matrix, but these target specific molecules. Untargeted approaches like FT-IR micro-spectroscopy would allow for a broader exploration. In this study FT-IR micro-spectroscopy analysis was implemented on sliced anaerobic granular sludge to explore the EPS distribution. Visualization of single wavenumber absorbance showed a higher polysaccharide content in the EPS at the granule perimeter, shifting to a higher protein concentration toward the centre. The boundary of this shift was approximately 150 µm from the surface, which was in accordance with the layer of fermentative bacteria described in literature. The complexity in the polymer composition meant that many functional groups were overlapping, making FT-IR annotation challenging. To address this, principal component analysis and two-dimensional correlation spectroscopy analysis were included in the analysis. These methods enabled the identification of overlapping functional groups and correlations between functional groups. Positive correlations between protein and polysaccharide functional groups suggested the presence of glycoproteins, which has been regularly described in chemical EPS analysis studies. Additionally, correlations between sulfated compounds and protein/polysaccharide functional groups indicated potential co-localization in the extracellular matrix. Differences in positive correlations of sialic acids with polysaccharides suggest variations in polysaccharide compositions, possibly caused by differences in the microbial community.

## 1. Introduction

Using granular sludge in wastewater treatment makes it possible to remove organic pollutants at higher loading rates compared to flocculent sludge (Van Lier et al., 2015). Microorganisms form granules by secreting polymers that embed them in an extracellular matrix (Flemming & Wingender, 2010). Extracellular polymeric substances (EPS) are a mixture of polymers found in the extracellular matrix, and in general contains: proteins, polysaccharides, lipids, extracellular DNA and combinations of these polymers (Flemming & Wingender, 2010; Seviour et al., 2019). EPS provide the structural support, and thus influence matrix stability (Felz et al., 2016; Bou-Sarkis et al., 2022). This structural support is caused by stabilization through either covalent bonding between polymers, hydrogen bonding or through cross-linking of charged groups in the EPS (Costa et al., 2018; Lotti et al., 2019). Microorganisms adapt to environmental cues by secreting charged moieties that influence the diffusion of ions into the extracellular matrix (de Graaff et al., 2019; Gagliano et al., 2018; Li et al., 2018). Amongst the most acidic charged moieties are sialic acids, phosphorylated and sulfated compounds. Examples of charged groups were found to be present in the EPS of many different biofilm systems (Boleij et al., 2020; de Bruin et al., 2022; Chen et al., 2023; Felz et al., 2020; Bourven et al., 2015). In both aerobic and anaerobic granules it was seen that sialic acids were more prevalent in high molecular weight fractions, whereas sulfated compounds were present throughout the whole molecular weight range (Chen et al., 2023). Whether this influences the functionality is not yet known, as the function of a specific polymer inside the extracellular matrix is also dependent on the location.

Seviour et al., (2019) proposed a roadmap for understanding the role of individual EPS components inside the extracellular matrix. Two routes have been suggested for imaging biofilms. The first is to extract structural EPS and determine the specific polymer structure, through chemical analysis and use this information to find specific probes for these chemical structures (Figure 1). Traditionally this is done by extracting the EPS from the biofilm using e.g. alkaline EPS extraction, and then analysing it using colorimetric and chemical analysis (Felz et al., 2019). Examples of these chemical analysis are nuclear magnetic resonance monomer analysis and high performance anion exchange chromatography with pulsed amperometric detection to identify polysaccharides and liquid chromatography combined with tandem mass spectrometry for protein identification (Gonzalez-Gil et al., 2015; Dubé and Guiot, 2019). The identified polymers can then be imaged by targeted imaging techniques like e.g. fluorescent lectin binding analysis for glycans and antibody for proteins (Neu & Kuhlicke, 2022). The advantage of this route is that specific compounds can be localized with a small optical resolution, making it possible to image the EPS of single cells inside the biofilm. The second route is by using untargeted imaging techniques like e.g. MALDI-TOF MS, Raman micro-spectroscopy and FT-IR micro-spectroscopy (Gowen et al., 2015). With these techniques, the spatially resolved spectra offer more chemical information than can be generated with typical microscopy. As such, these techniques detect a broad range of polymers that are measured simultaneously. The function of a specific polymer relies on the presence of co-localized polymers/monomers. Thus, in comparison to the targeted imaging route, the untargeted imaging route provides broader imaging capabilities.

**Figure 1.**
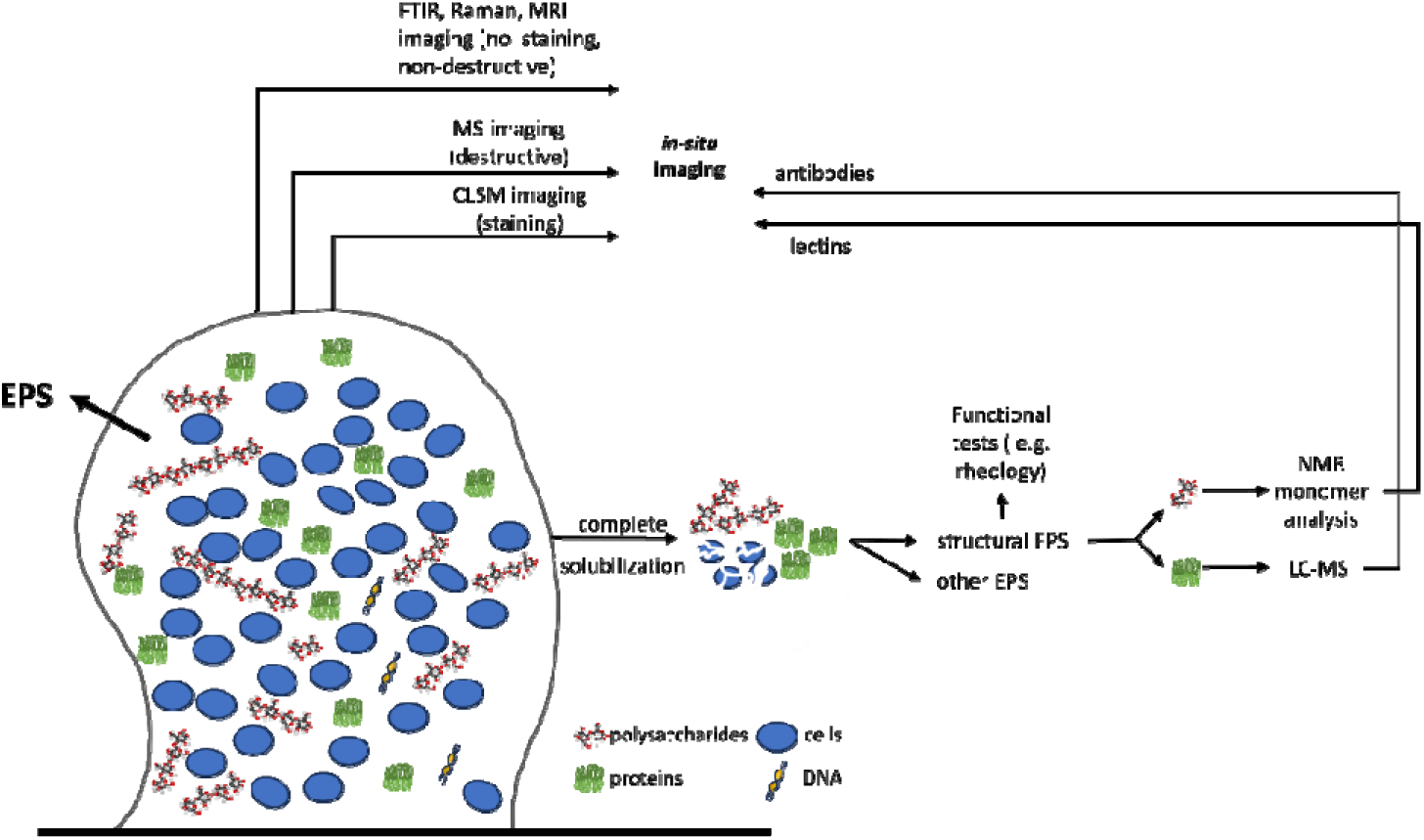
Roadmap adapted from Seviour, et al. (2019), showcasing potential routes for biofilm imaging.

With FT-IR technique, many different functional groups present in polymers can be detected because vibrational spectra are molecule specific. FT-IR imaging is a user-friendly and non-destructive technique with minimal sample preparation and a high signal to noise ratio (Gowen et al., 2015). Mapping of the polymer distributions can be achieved by using the absorbance height, or area, of bands associated with specific polymer types (Jiang & Rieppo, 2006). However, absorbance bands of functional groups often correspond to multiple polymer types, leading to spectral complexity. In these cases, it is beneficial to utilize all the information stored in the spectrum simultaneously. This task is commonly addressed with chemometrics, a field that applies mathematical and statistical methods to analyse chemical data. A simple way of reducing the complexity of the spectral data is by reducing the dimensionality. With principal component analysis (PCA) this data dimensionality reduction is achieved while retaining the majority of the information. PCA describes the absorbance spectra as a linear combination of component spectra (principal components) and their scores. By imaging the principal component scores, it is possible to visualize the polymer distribution with limited loss of information (Gowen et al., 2015; Chirman & Pleshko, 2021). To see if polymers co-localize in the extracellular matrix the corelation of different functional groups can be explored. Generalized two-dimensional correlation spectroscopy (2D-COS) is a way to analyse correlations in the spectra between absorbance bands under the influence of e.g. time, pH, temperature or space (Jiang & Rieppo, 2006; Noda et al., 2000). This technique is used to determine the correlation spectrum of spatially resolved absorbance spectra (Lasch & Noda, 2019). If the functional groups are correlated, it indicates those polymers are co-localized.

The aim of this study is to explore the extracellular matrix polymer composition using FT-IR micro-spectroscopy. Through the use of chemometrics a more guided effort can be achieved of the untargeted approach to visualize the polymer composition in the extracellular matrix. The use of FT-IR micro-spectroscopy may enhance our understanding of the spatial arrangement of polymers in the granular sludge matrix, providing valuable insights into its composition and structure.

## 2. Materials and methods

### 2.1. Sample handling and FT-IR screening of EPS absorbance bands

Granular sludge was collected from a full scale anaerobic wastewater treatment plant treating papermill effluent similar to other studies before (Chen et al., 2023; Doloman et al., 2024). The first step in the analysis of the extracellular matrix is to identify the functional groups present in the FT-IR absorbance spectrum of EPS. EPS extraction was performed on washed and lyophilized granular sludge using an alkaline extraction method as described earlier (Pinel et al., 2020). FT-IR spectra of the extracted EPS were recorded with a FT-IR spectrophotometer (Spectrum 100, PerkinElmer, Shelton, USA) over a wavenumber range of 4000 cm^-1^ to 600 cm^-1^ with 16 accumulations and 4 cm^-1^ spectral resolution.

### 2.2. Homogeneous granule slicing and data acquisition

FT-IR micro-spectroscopy using an ATR-imaging crystal is a technique that measures the absorbance spectra at the surface of the ATR crystal. Therefore, if unsliced granular sludge is used for the imaging analysis, the external surface can be imaged. Slicing of the granular sludge ensures the possibility of imaging the internal granular structure. Before slicing, granules of 1 mm in diameter were selected and were embedded in an optimal cutting temperature medium (Neg-50™ Frozen section media, Epredia™). Granules were sliced with a cryotome to obtain slices that were 15 µm in depth (cabinet temperature -20°C, mount temperature -25°C; Leica CM1900).

FT-IR micro-spectroscopy imaging was performed in ATR imaging mode with a germanium attenuated total reflectance crystal (Spotlight 400 FTIR microscope, PerkinElmer, Shelton, USA). Absorbances were recorded in the 4000 cm^-1^ to 600 cm^-1^ range with 4 accumulations, a spatial resolution of 1.56 µm^2^ and a spectral resolution of 4 cm^-1^. The measured area could be set to be anywhere between 50 x 50 µm and 500 x 500 µm, and was set to 300 x 300 µm for this study.

### 2.3. Data visualization

The workflow for FT-IR imaging as employed in this study is shown in figure 2. The steps are discussed in detail in the following text, for additional information how this is applied see the supplemental MATLAB script (Supplemental information S1).

**Figure 2.**
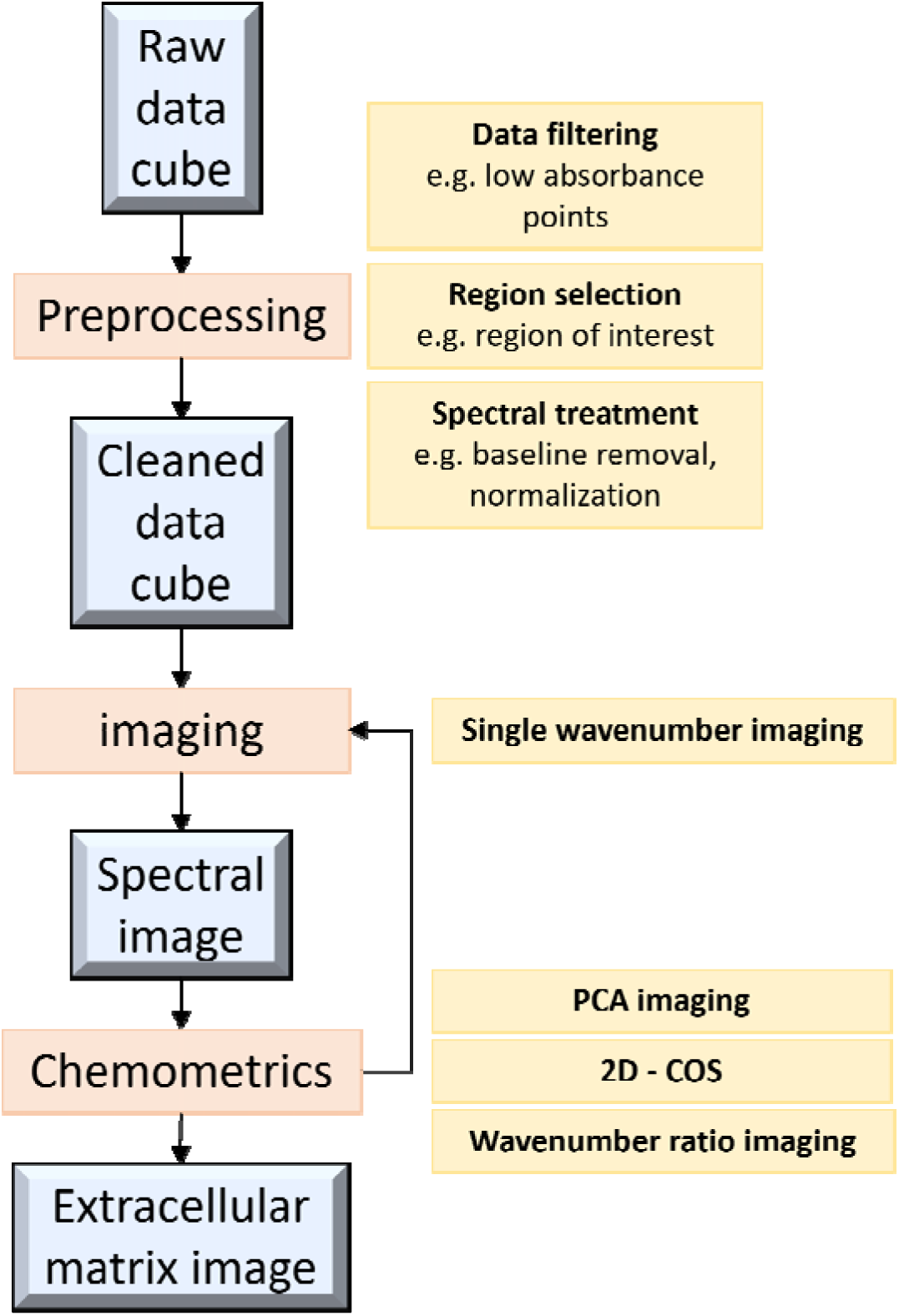
workflow diagram FT-IR imaging data pipeline.

#### 2.3.1. Data pre-processing

For the acquisition of the data it was important to ensure good sample contact between the sample and imaging crystal. This would make it possible to obtain data with a high signal over noise ratio. After acquisition the data was smoothed with a Savitsky-golay filter, and base-line was corrected by a zero-order baseline correction. Subsequently normalization was performed by feature scaling. Due to the heterogeneity of the biofilm, there are discrepancies on the average absorbance height between locations on the granule slice. This can partly be solved by ensuring slicing as homogeneously as possible. Supplemental Figures S1-S2 show the microscopic images of the sliced granular sludge. The grooves of the metal disc on which the samples were mounted affect the absorbance height (Supplemental Figure S4). The average absorbance height images were used to locate and filter areas with a high signal to noise ratio. Based on the data, pixels with an average absorbance less than 0.3 were discarded (Supplemental Figures S3-S4).

#### 2.3.2. Single absorbance band imaging

The single point absorbance imaging was created by mapping the absorbance height of proteins (Amide I band at 1630 cm^-1^), polysaccharides (C-O-C band at 1030 cm^-1^), lipids (C–H stretch of >LCH_2_ groups, dominated by lipid contributions at 2920 cm^-1^) (Talari et al., 2017), sialic acids (CL=LO stretch of ester specifically correlated to sialic acids at 1726 cm^-1^) (Nallala et al., 2020) and putative sulfate groups (sulfate stretch at 1206 cm^-1^) (Brézillon et al., 2014). The absorbance height for all the given wavenumbers was then scaled and mapped across the measured area to show the relative differences in distribution.

#### 2.3.3. Principal component analysis

The interpretation of large datasets, like those from FT-IR imaging measurements, is challenging. Interpretation of data is increasingly complex in FT-IR absorbance spectra due to the presence of broad and convoluted absorbance bands. To address this, PCA is employed, which reduces highly dimensional data to a more manageable format, while conserving as much of the variability of the data as possible. The application of PCA results in the deconstruction of the absorbance signal into two matrices: the loadings matrix and the scoring matrix. The loadings matrix assigns weights to each wavenumber for every principal component, while the scoring matrix assigns scores to each absorbance spectrum for every principal component. The first principal component captures the highest amount of variability. Subsequent principal components are independent of each other, each describing the most variability in turn. In this study, the PCA loadings and scores were calculated using the built-in function in MATLAB. This methodology enables a more accessible and insightful understanding of the intricate FT-IR data, facilitating the extraction of meaningful patterns and information from the complex spectra (Jolliffe & Cadima, 2016).

#### 2.3.4. Two-dimensional correlation spectroscopy

Generalized two-dimensional correlation spectroscopy is a powerful analytical method for the interpretation of vibrational spectra that is known primarily from use in NMR spectroscopy. Its use however, is not limited to NMR and can be used in other vibrational spectroscopy techniques as well (Noda et al., 2000). Variations in the spectrum arising from, in this case, spatial differences can serve as input for the correlational analysis. The synchronous 2D correlation spectrum essentially mirrors the covariance matrix. The matrix is derived by calculating the covariance between all combinations of spectral feature vectors at the specified spectral variables v_1_ and v_2_. By applying color-coding to represent the covariance values at v_1_ and v_2_ a 2D covariance map can be generated (Lasch & Noda, 2019).This map visually encapsulates the interrelationships among spectral features and provides an overview of covariations across FT-IR spectra.

## 3. Results

### 3.1. Determination of the FT-IR absorbance regions of interest

An initial screening of the functional groups present in EPS was performed using FT-IR on EPS. To get an understanding of what absorbance bands are important in EPS, first an extracted EPS sample was analysed. The EPS appears to be protein dominant as shown in the height of the protein region absorbance bands. The relatively high and wide bands in the polysaccharide region indicate a high polysaccharide content that are likely present in large polymer chains. Other functional groups include absorbance bands annotated to the lipid region and anionic compounds like phosphate, sulfate and sialic acids (Figure 3).

**Figure 3.**
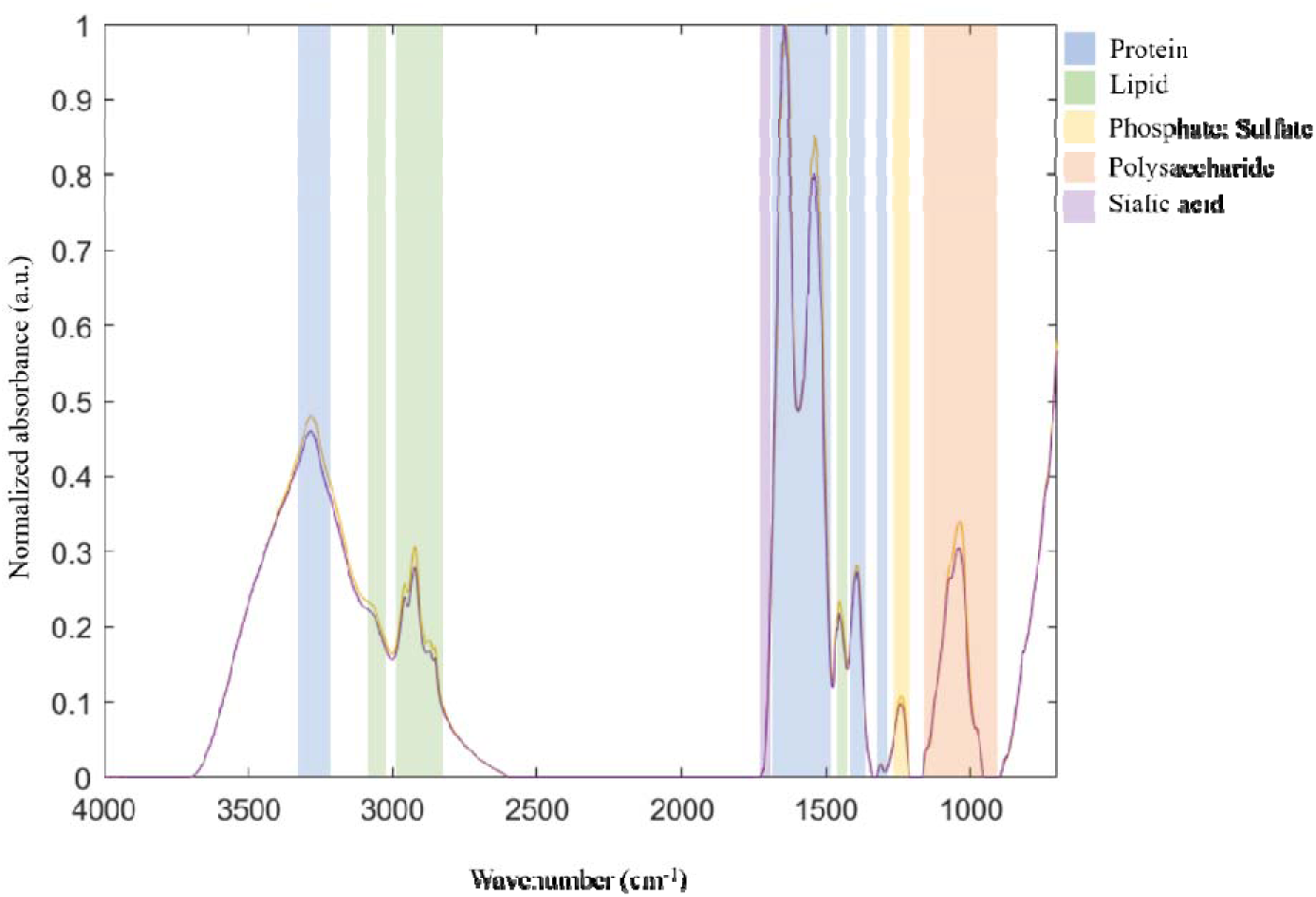
FT-IR absorbance spectra of EPS extracted from anaerobic granular sludge with annotated polymer functional region absorbance bands.

### 3.2. Single-point absorbance

The chemical distribution of the most prevalent components in EPS were mapped (Figure 4) for a sliced granule. Firstly, it can be seen that, all the polymers and sialic acids are heterogeneously distributed in the granule. For example, in the protein chemical image (figure 4A), it can be seen that the bottom left corner is more protein rich relative to other mapped areas (indicated by arrow). Similarly, there are specific areas where polysaccharides, lipids, sialic acids and sulfated/phosphorylated polymers are more abundant than at other areas (indicated by the arrows at the specific chemical image). Secondly, there are both positive and negative correlations between the signals of different components i.e. polysaccharides and proteins signals are in general positively correlated except for the top and top right zone. The lipid rich regions were distributed similarly to the protein rich regions. Sulfated/Phosphorylated polymers followed similar abundancy profiles as polysaccharides and lipids (as indicated in 4B and 4C) except for the area indicated by the arrow in 4E. In comparison, the sialic acids signal seemed to be negatively correlated with the signal of other components.

**Figure 4.**
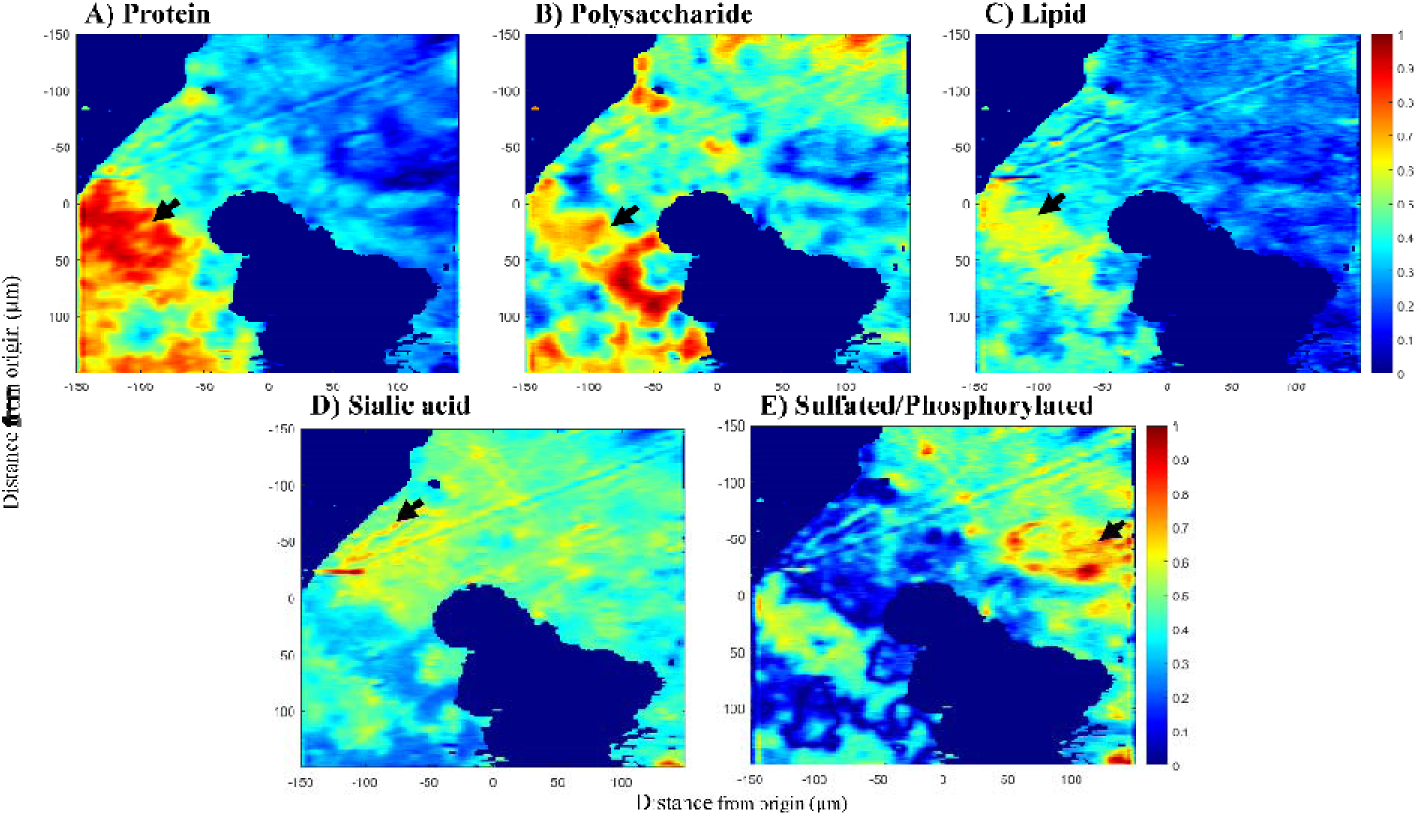
Scaled mapping of FT-IR functional groups absorbance band heights of sliced anaerobic granular sludge treating papermill wastewater. Protein) Height of the protein region absorbance band at 1630 cm^-1^. Polysaccharide) Height of the polysaccharide region absorbance band at 1030 cm^-1^. Lipid) Height of the lipid region absorbance band at 2920 cm^-1^. Sialic acid) Height of the sialic acid region absorbance band at 1726 cm^-1^. Sulfate) Height of the putative sulfate region absorbance band at 1206 cm^-1^. The origin in this picture is the centre of the ATR imaging crystal, and so also the centre of the imaged area. Regions that were filtered out due to weak signal are shown in dark blue. The black arrows indicate regions of high absorbance.

### 3.3. Principal component analysis for dimension reduction

Proteins, polysaccharides, lipids and sialic acids contain multiple functional groups. The signal of these functional groups can overlap with each other which makes it difficult to get a complete characterization by only looking at single wavenumber absorbance bands for each component. In order to visualize the biggest differences in the data set with the complex combinations of functional groups whilst still maintaining most of the information at a reasonable level, principal component analysis (PCA) was applied. Figure 5 shows the spectrum of the second, third and fourth principal component (PC) loadings calculated from the spectral data. While the first PC could explain most of the variability in the area (67.9%), it showed the biggest variety in the data could be explained by the absorbance intensity of most functional groups. To image how more specific compounds were distributed across the granule slice, the second, third and fourth PC were chosen. These explained 23.6%, 3.6% and 1.3% of the variability in the data, respectively. The loading of PC2 was protein dominant and emphasized amide functional groups: 3300, 3070, 1660, 1560, 1500 and 1240 cm^-1^. However, other functional groups like lipid CH_3_: 2960 and 2900 cm^-1^, polysaccharides: 1110, 1070 and 1000 cm^-1^ and sialic acid at 1728 cm^-1^, where also present in the PC2 loading. The PC3 loading was more polysaccharide dominant and emphasized polysaccharide functional groups:1300, 1132 and 968 cm^-1^. Other important functional groups were lipid CH_2_ stretch: 2994 and 2832, sulfate/phosphate:1218 cm^-1^ and amide functional groups: 1676 and 1480 cm^-1^. Lastly, PC4 was polysaccharide dominant with highest weight on the polysaccharide functional groups: 1100 and 1014 cm^-1.^ Additional functional groups were amide: 1684, 1668 and 1580 cm^-1^, and CH_2_ stretch 2920 cm^-1^.

**Figure 5.**
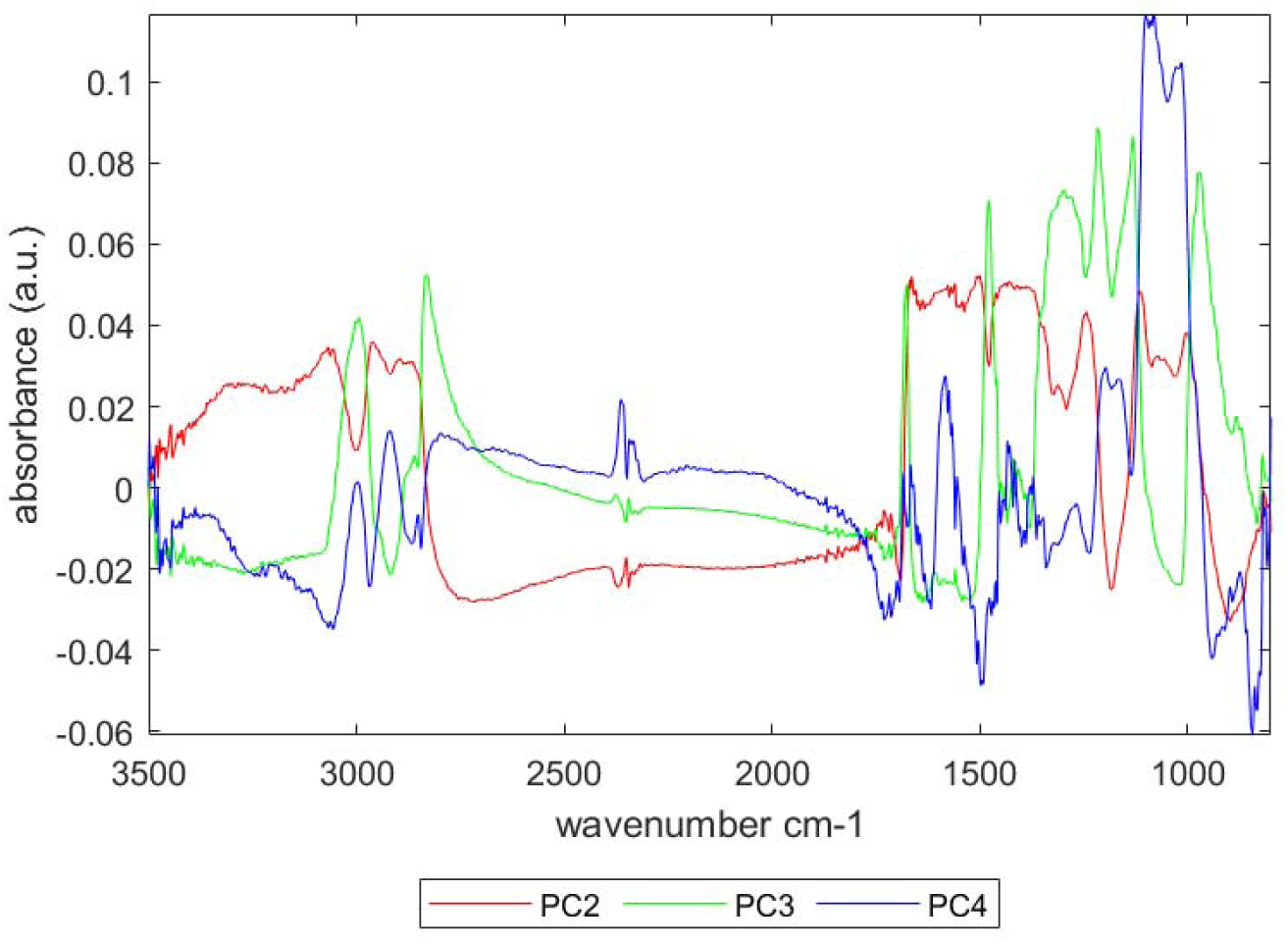
Principal components loading spectrum of a granular sludge slice.

The PC scores of the spatially resolved FT-IR spectra were mapped and are shown in figure 6. The left bottom part had a relatively high score for PC2, which coincided with the mappings of the protein and part of the polysaccharide, lipid and sulfate functional groups seen in figure 4. The region which the highest overlap of functional groups was marked in yellow indicating an overlap of PC2 and PC3 scores. Scores on PC3 were also high in the right region of the mapped area. This region had relatively high absorbance for sulfated/phosphorylated functional groups. High scores on PC4 indicated a more dominant presence of polysaccharides and the regions with high scores on PC4 also had relatively higher polysaccharide functional group absorbance in these areas. Mapping of the PC scores showed that there are many overlapping functional groups, but that it was possible to distinguish between similar functional groups by looking functional group pairs found in the PCs.

**Figure 6.**
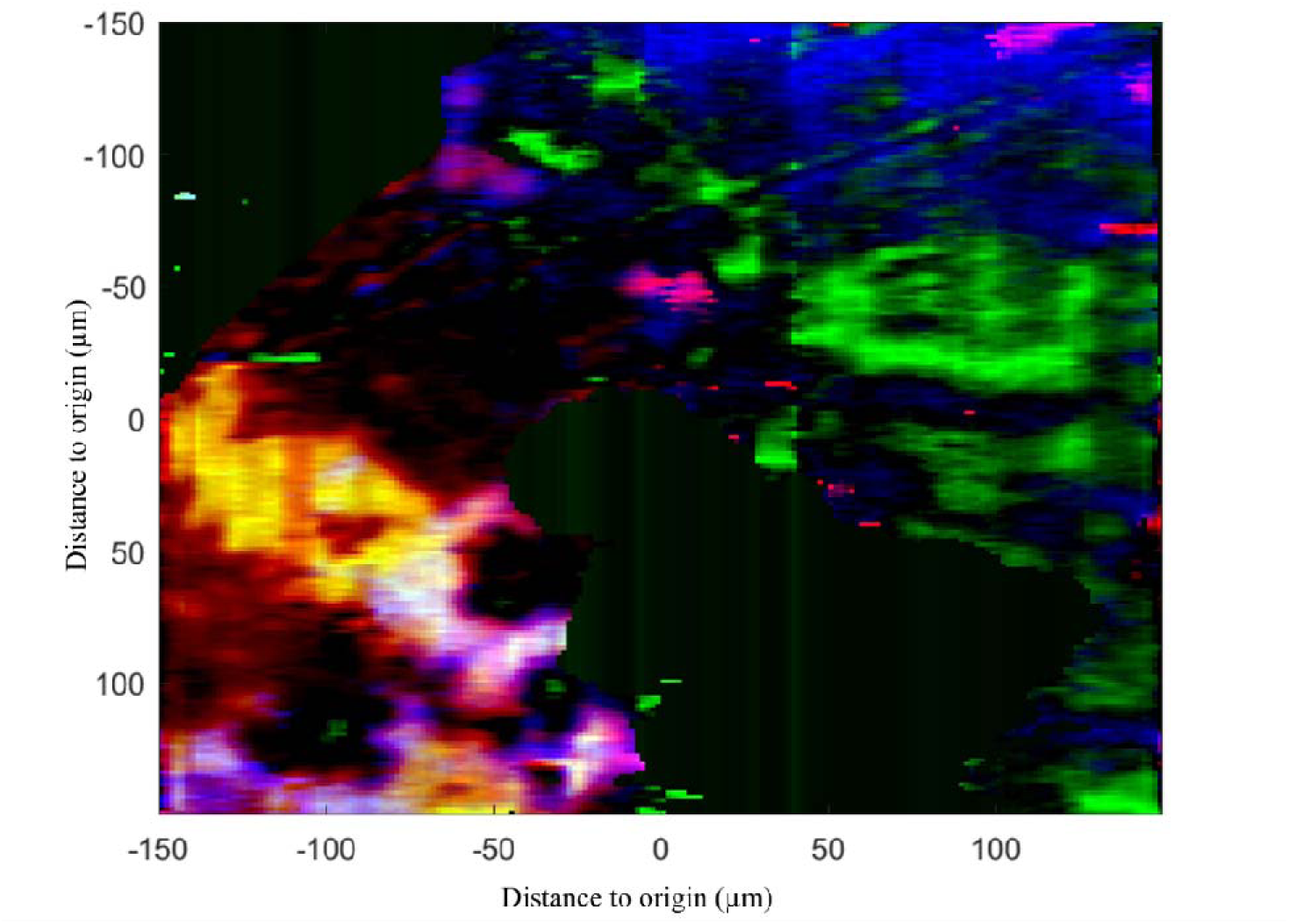
Mapping of the PC scores for PC2 (red), PC3 (green) and PC4 (blue) across the granule slice.

### 3.4. 2D-COS analysis

The EPS composition in anaerobic granules was spatially heterogeneous and there was strong overlap of bands from different components. In order to decipher the correlations between overlapping functional groups and detect compositional changes which are ‘hidden’ from PCA and single wavenumber absorbance, two-dimensional correlation spectroscopy (2D-COS) analysis was used. The spatially resolved FT-IR absorbance spectra were used to generate the synchronous correlation spectrum shown in figure 7. It is observed that, for each individual component, there are positive correlations among their specific typical absorbance regions along the scanned surface area. i.e. –For polysaccharides, absorbance at the regions of 850 – 920 cm^-1^, 1000 – 1050 cm^-1^ and 1160 – 1200 cm^-1^ are positively correlated. While for proteins, amide regions at 1380 – 1460 cm^-1^ and 1500 – 1700 cm^-1^ were also positively correlated. The functional group annotated to sialic acid (1726 cm^-1^) was also positively correlated along the scanned area.. For lipids, the only functional group that was positively correlated was for the CH_3_ stretch found at 2840 – 2880 cm^-1^. Secondly, in between different components, positive correlations were observed among lipid, polysaccharides and proteins at 2960 and 2860 cm^-1^ (-CH stretch, with lipid dominantly contributed), 1000 – 1050 cm^-1^ (C-O-C stretch) and 1380 – 1460 cm^-1^ and 1500 – 1700 cm^-1^(amide bands). Interestingly, lipid seemed to be positively correlated with the sialic acid functional group (at 1726 cm^-1^) only at 2920 cm^-1^ and not at other regions annotated to lipids. The sialic acid functional group was mainly correlated with the polysaccharide functional groups at 850 – 920 cm^-1^, 1000 – 1050 cm^-1^ and 1160 – 1200 cm^-1^ regions. Proteins and polysaccharides were positively correlated at 1630 cm^-1^ and the 1000 – 1050 cm^-1^ spectral regions. Phosphorylated or sulfated polymers at 1234 cm^-1^ seemed to be mainly positively correlated with the polysaccharide and protein functional groups at 1000 – 1050 cm^-1^ and 1500 – 1700 cm^-1^, respectively. Another spectral band appeared at 1180 cm^-1^ was observed positively correlated with the sialic acid band at 1726 cm^-1^, but not with any other absorbance.

**Figure 7.**
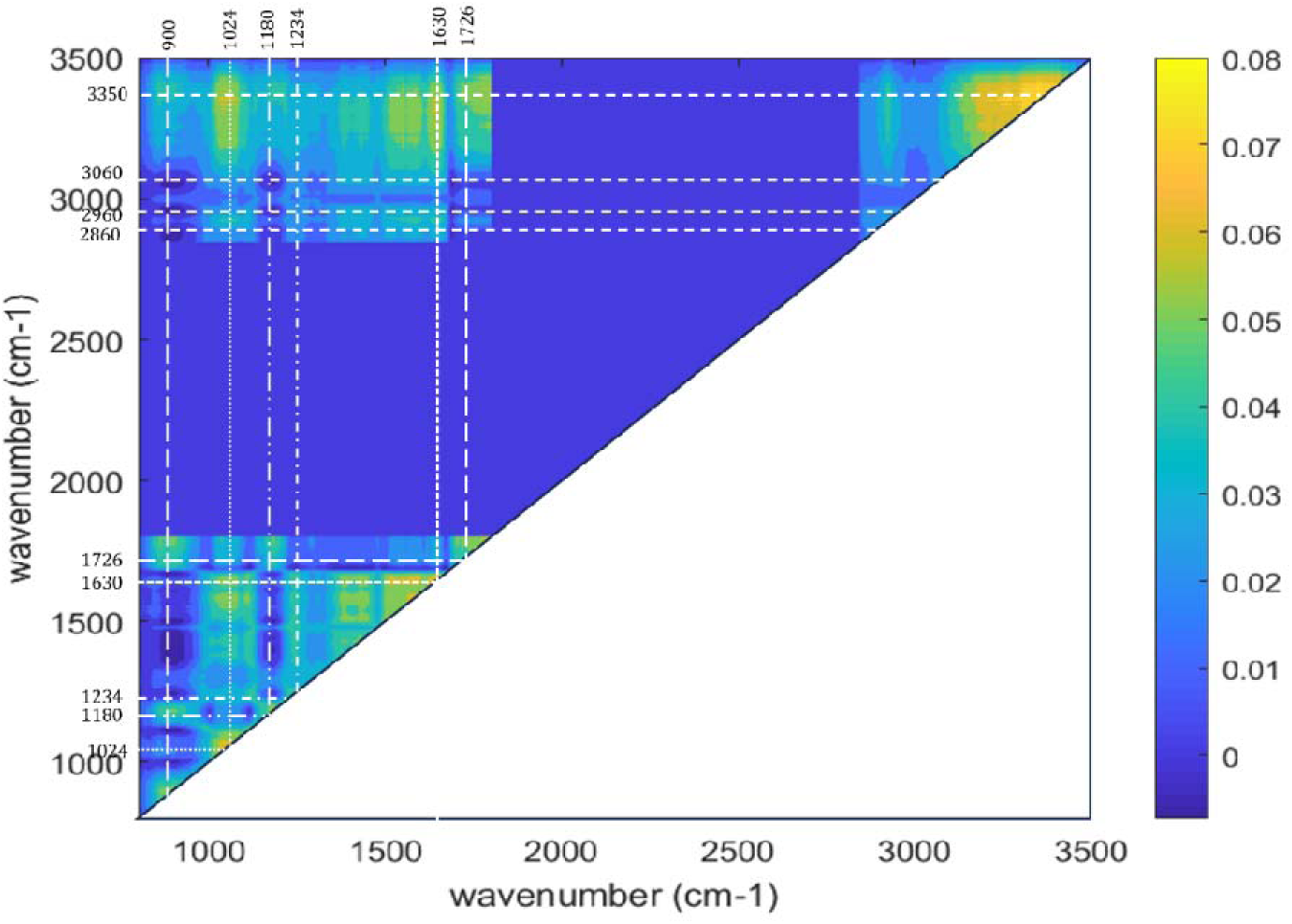
Generalized synchronous two-dimensional correlation spectrum contour plot derived from the spatially resolved spectra.

### 3.5. Imaging at the outer edge of the anaerobic granular sludge

To see if the general reported layered structure in anaerobic granules could be found in the granular sludge closer to the outside of the granule, a sample from the same WWTP was imaged at the edge of the granule, i.e. close to the surface of the granule towards the external environment (Sekiguchi et al., 1999; Satoh et al., 2007). The protein content was distributed unevenly in the mapped area. High absorbance height was found when going more towards the centre of the granule, as indicated by the white arrow. In contrast, polysaccharides were found to have a higher absorbance height in closer towards the outer edge of the measured area. Lipid absorbance height had great similarity with the sialic acid, and protein distribution. With the lipid and sialic acid overlapping mostly in the region indicated with the white arrow in the sialic acid mapping. Sulfated compounds also was similar to the sialic acid and protein distribution but differed from sialic acids with a higher absorbance at the white arrow. The protein dominant region (indicated by the arrow), had a higher absorbance for the sulfated compounds than for the sialic acid compounds (figure 8).

**Figure 8.**
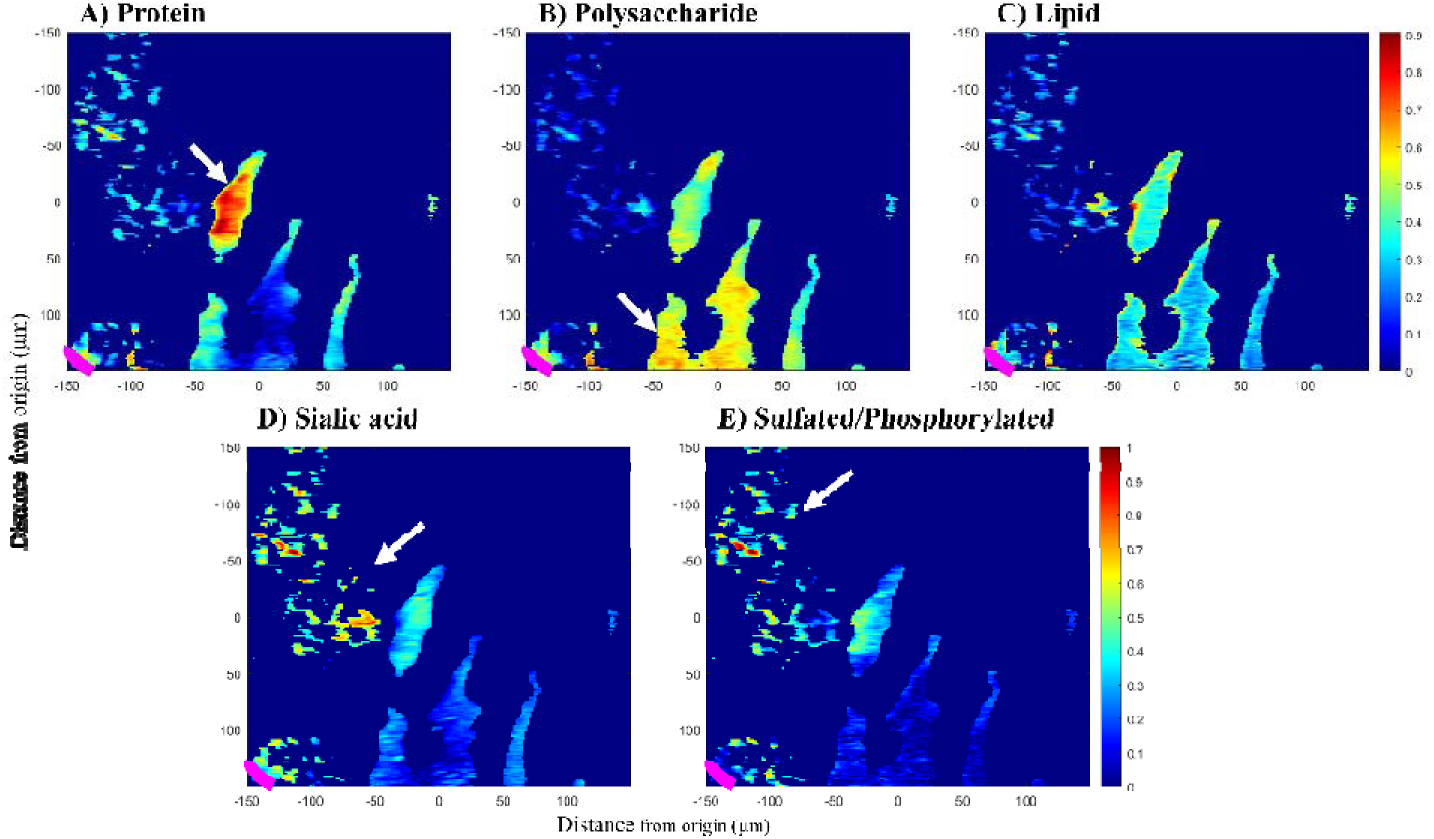
Scaled mapping of FT-IR functional groups absorbance band heights of sliced anaerobic granular sludge treating papermill wastewater imaged close to the edge of the granule. A) Height of the protein region absorbance band at 1630 cm-1. B) Height of the polysaccharide region absorbance band at 1030 cm-1. C) Height of the lipid region absorbance band at 2920 cm-1. D) Height of the sialic acid region absorbance band at 1726 cm-1. E) Height of the putative sulfate region absorbance band at 1206 cm-1. The origin in this picture is the centre of the ATR imaging crystal, and so also the centre of the imaged area. Regions that were filtered out due to weak signal are shown in dark blue. The white arrows indicate regions of high absorbance. The pink line indicates the outer edge of the granule.

#### 3.5.1. Chemometric analysis on area on the outer edge of the granule

Figure 9 shows the spectrum of the second, third and fourth PC loading spectra. Like the PCA performed on the centre part of the granule, the first principal component could explain the highest variability of the data (86.6%). The biggest variety was mostly caused by the presence of almost all functional groups. To image more specific functional groups, imaging was performed using the second, third and fourth PC (5.7%, 3.8%, 1.0%), respectively. Functional groups contributing to the second biggest variety of the data as seen in PC2 mostly polysaccharide in nature: 3300, 1084 and 1034 cm^-1^. In PC2 protein absorbance bands were actually negatively contributing and composed of wavenumbers: 3060, 1676, 1580 and 1476 cm^-1^. In contrast, PC3 was mainly protein dominated with absorbance bands at: 1620, 1530, 1450 and 1380 cm^-1^. Other functional groups of interest where lipids found at absorbance bands: 2960 and 2870 cm^-1^, and polysaccharides at: 1144 and 956 cm^-1^. In PC4, protein absorbance bands were the main contributors in the wavenumbers: 1636 and 1540 cm^-1^. Additionally, sialic acid absorbance bands at 1726 cm^-1^ contributed to PC4. Sulfate and polysaccharide were negatively contributing with the following wavenumbers: 1246, 1110, 948 and 854 cm^-1^, respectively.

**Figure 9.**
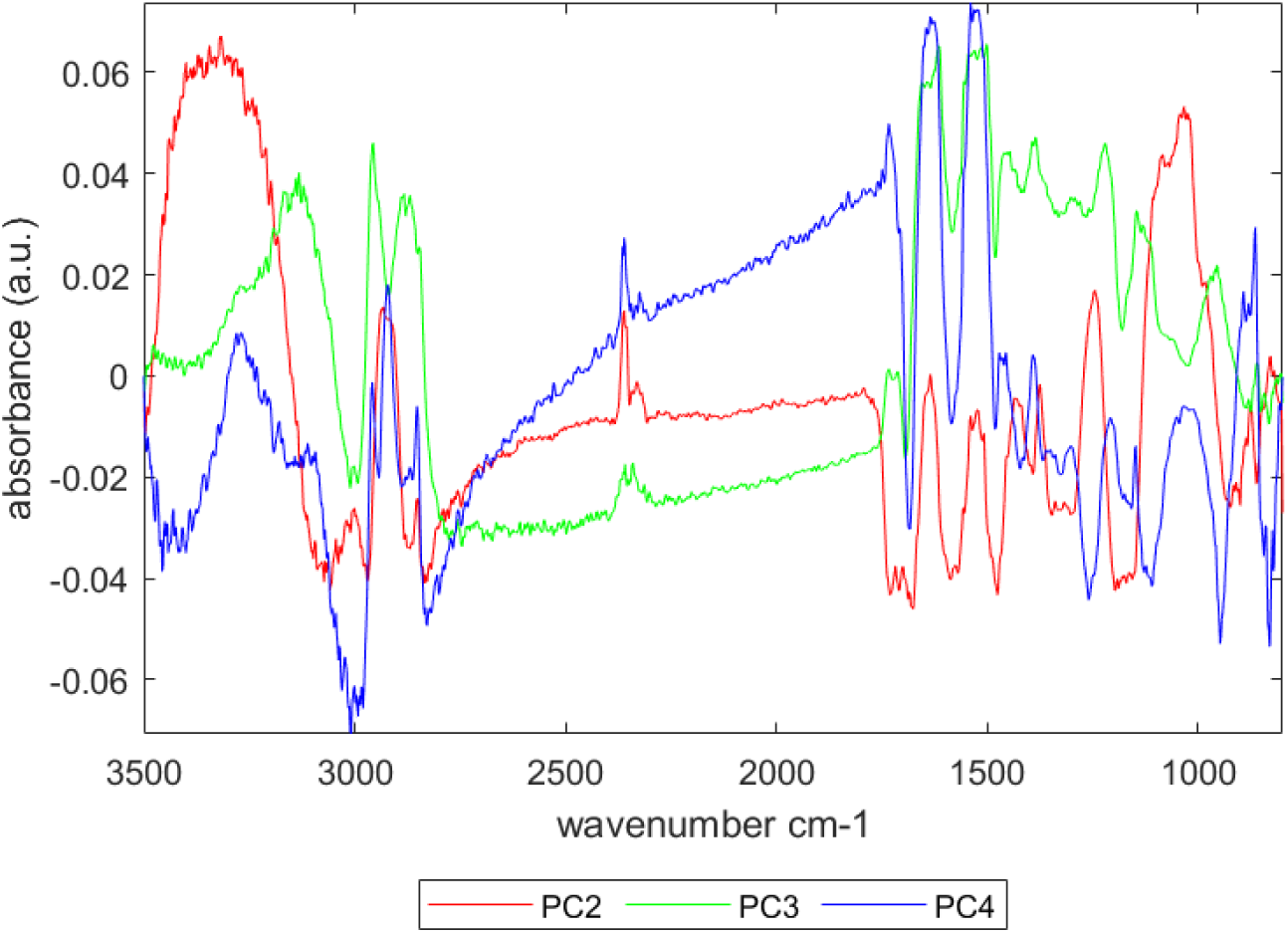
Principal components loading spectrum granular sludge slice at the outer edge.

The scoring for PC2, PC3 and PC4 of the spatially resolved spectra were mapped and are shown in figure 10. In the region close to the outer edge of the granule, high scores for PC2 (red) indicated a relatively higher presence of polysaccharides and sulfated polymers. Going toward the centre of the granule PC3 (green) and PC4 (blue) were more prevalent. High scores on PC3 and PC4 both indicated a protein dominant spectrum, but PC3 also contained absorbance for sulfated polymers and polysaccharide absorbance bands at 1154 and 956 cm^-1^. Whereas, PC4 had a negative absorbance in the above mentioned polysaccharide bands, with a high absorbance at 854 cm^-1^. The majority of the spectra scoring high on protein dominated PC loading spectra, were found towards the centre of the granule. Conversely, spectra scoring high on polysaccharides were found primarily near the edge of the granular sludge.

**Figure 10.**
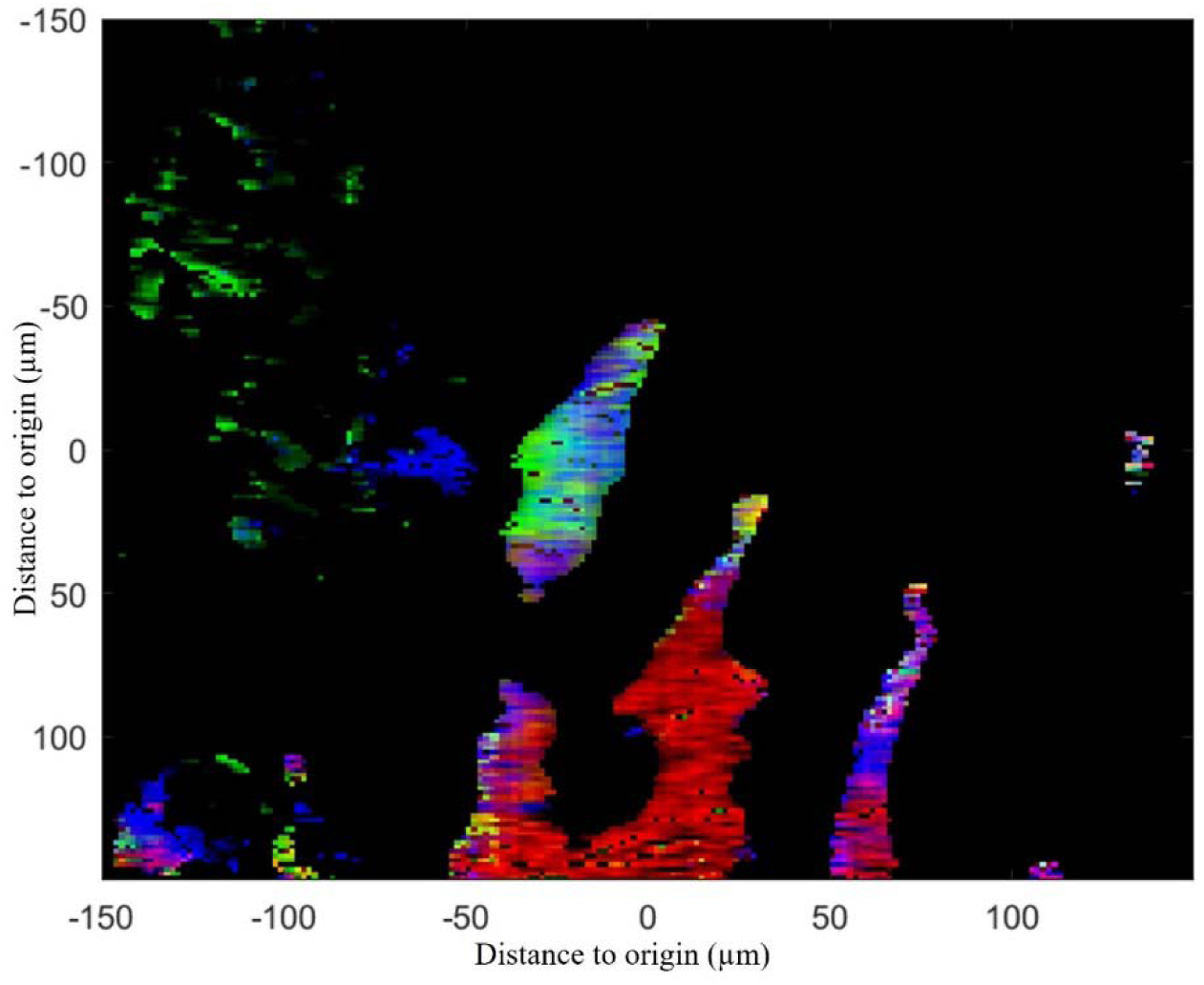
Mapping of the PC scores for PC2 (red), PC3 (green) and PC4 (blue) across the granule slice at the outer edge of the granule.

#### 3.5.2. 2D-COS spectrum of the FT-IR spectra from the outer edge of the granule

The 2-D correlation spectrum for the FT-IR spectra is shown in figure 11. The most prominent functional groups visible in the data were sialic acid (1726 cm^-1^), amide I & II (1630 & 1540 cm^-1^), sulfated compounds (1234 cm^-1^) and polysaccharides (1100 and 1024 cm^-1^). The correlation spectrum on the area near the granule edge was similar to the correlation spectrum made from the data in the centre of the granule shown in figure 7.

**Figure 11.**
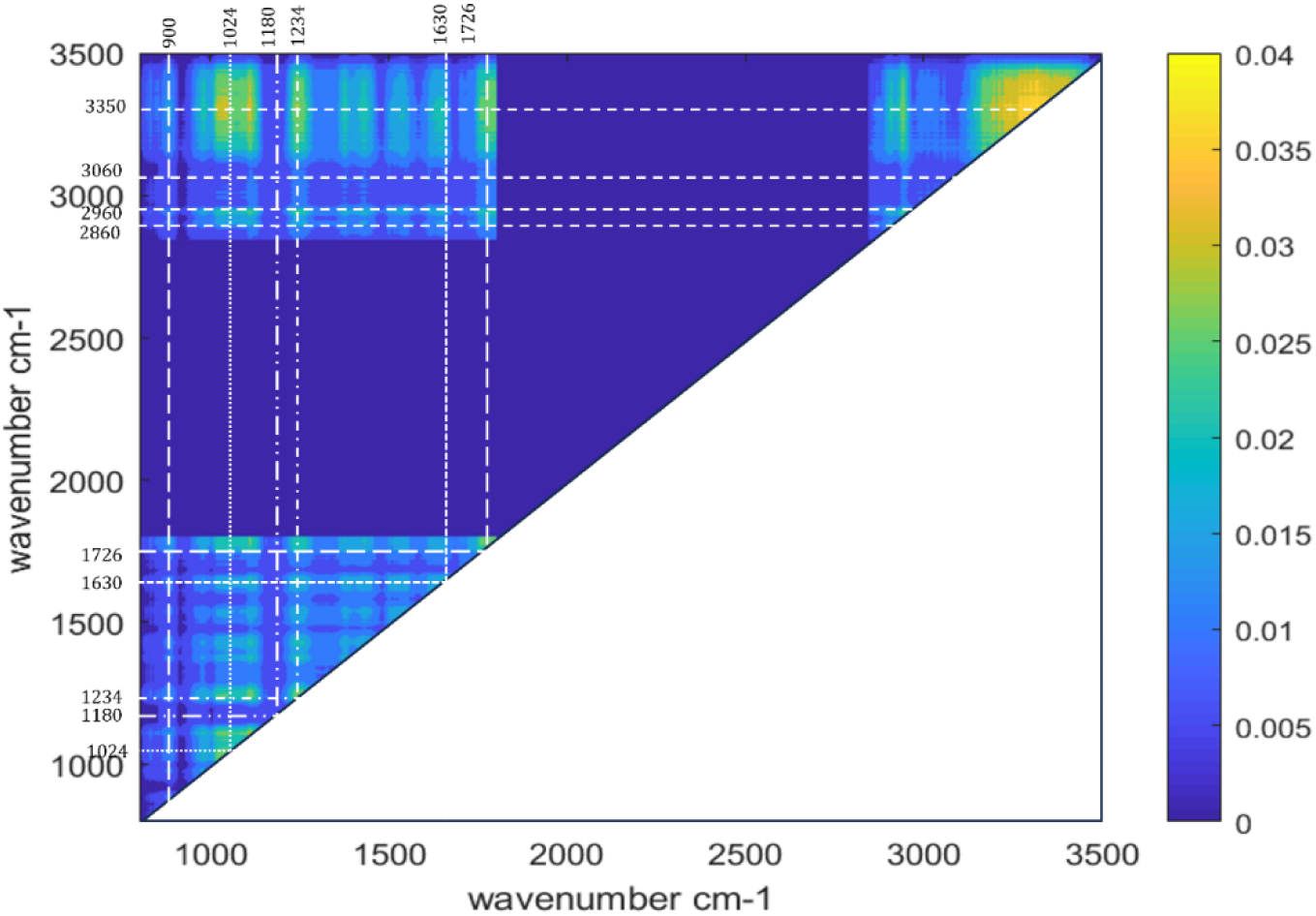
Mapping of the PC scores for PC2 (red), PC3 (green) and PC4 (blue) across the granule slice at the outer edge of the granule.

## 4. Discussion

### 4.1. The spatial structure of extracellular polymeric substances revealed by FT-IR microscopy

In this study, the possible application of FT-IR micro-spectroscopy for the visualization of extracellular polymer distribution across the granular sludge was explored. Anaerobic granular sludge has been described to have a layered structure. Fermentative bacteria grow on the outside, with syntrophic and hydrogenotrophic bacteria further in the middle and acetoclastic methanogens at the center of the granule (Satoh et al., 2007). In the center, the EPS is protein dominated (Ding et al., 2015; Forster & Quarmby, 1995). However, this layered structure is not always the case and clustered structures have also been reported (Gonzalez-Gil et al., 2001). It should be noted that the fermentative and syntropic layer in the layered structure can vary in size but also depends on the heterogeneity of the samples. In the current research, it was observed that there are regions with different EPS content which might follow the layered structure. While as far as it is close to the center of the granules, a more clustered structure can be distinguished. There seems to be no correlation between distance from the center and polymer composition. The granule sludge diameter was around 1.6 mm, estimated from the measured area size the covered area is 4.5 % of the total granular sludge slice. This mapped area would give an accurate representation of the sample if the EPS components were heterogeneously distributed. However, when looking at the data at the outer edge of the granules, a higher polysaccharide abundance at the surface and a higher protein abundance next to it towards the center of the granule is observed. The boundary between the polysaccharide rich and the protein rich layers is about 150 µm from the granule surface, which is in accordance with other studies (Sekiguchi et al., 1999; Doloman et al., 2017). It is highly likely that the clustered EPS composition is caused by clustered communities of different microorganisms. The outside of the granular sludge seems to be dominated by a microbial community that produces a polysaccharide rich EPS, which could be attributed to fast-growing fermentative bacteria, which can easily obtain the substrate from the environment. Below the fermentative layer, the EPS composition did seem to be more randomly distributed and more clustered than layered.

### 4.2. Correlations in extracellular polymer composition unveiled by chemometric analysis

There were regions in the granular sludge slice that showed strong overlapping of functional groups in the EPS. This is partly caused by the convoluted absorbance spectra in biological structures (Marcott et al., 2014). Biological samples consist of complex polymers which are covalently linked proteins, polysaccharides, lipids and other molecules such as glycoproteins and glycolipids (Seviour et al., 2019). When covalently linked, they would co-localize in the spatial distribution. However, this is not the only reason why multiple absorbance bands would overlap in the spatial distribution map. Even a single polymer without covalent bonding has multiple functional groups that will generate multiple absorbance bands in varying degrees. By evaluating the increased or decreased absorbance in combinations with absorbance bands, it is possible to find features that are otherwise ‘hidden’ in the spectra (Jiang & Rieppo, 2006; Marcott et al., 2014; Ali et al., 2018). This is the basics of chemometrics, which focuses on exploratory data analysis that provides an overall view of the biological samples under study, allowing to catch possible similarities/dissimilarities among samples, to identify the presence of clusters or, in general, systematic trends, to discover which variables are relevant to describe the sample. The absorbance height figures of the single absorbance bands (figures 4 and 8) could visualize the presence of polysaccharides. However with the figures showing the PCA (figures 6 and 10), it could be seen which other functional groups would also be found present at the same place. Especially in figure 4, there are two regions that contain a high polysaccharide peak absorbance. In the PCA figure it is shown that one region scores high on PC2 and PC3, whereas the other region scored higher on PC4. By looking at the PCA scoring images (figures 6 and 10), it is observed that even though high polysaccharide regions were found at multiple locations, there can be different compositions in these regions. This distinction could be seen with PCA imaging. Therefore, chemometric techniques make it possible to look at multiple absorbance bands at the same time and process possible correlations between absorbance bands. With 2D correlation spectroscopy, it was observed that the functional groups of proteins and polysaccharides were positively correlated, indicating a possible presence of glycoproteins. In fact, these covalently linked polymers have been identified in many studies targeting granular sludge (Boleij et al., 2020; Chen et al., 2023; Felz et al., 2020). Sulfated compounds were also shown to be correlated with protein and polysaccharide functional groups, indicating a potential co-localization in the extracellular matrix. The generated 2D-COS spectra of the different mapped areas were highly similar. However, differences could be observed in the positive correlations of sialic acids with polysaccharides. Figure 7 showed that sialic acids were positively correlated with polysaccharides and, in lesser extent, to proteins. Absorbance bands at 900 cm^-1^ and 1180 cm^-1^ were correlated to sialic acid absorbance bands. This correlation was not visible in figure 11, where a positive correlation was seen between sialic acid and absorbance bands at 900 and 1024 cm^-1^. This could indicate that the sialic acids are linked to different sugar monomers or even completely different polymers. In Chen et al., (2023) it was shown that in the largest molecular weight samples, the EPS was more glycosylated and contained the most sialic acids. From the 2D-COS data this seems to be the case as well, whereas sulfated polymers are correlated to all different types of polymers.

### 4.3. Data analysis and limitations

The data acquired from the FT-IR micro-spectroscope is in the form of a hyperdimensional data cube. The FTIR images were acquired on the sliced granules across a 300 μm x 300 μm area, with a pixel size of 1.56 μm x 1.56 μm, which results in a total of 36,864 pixels. Within each pixel, the FT-IR absorbance spectrum was recorded across the 4000 – 600 cm-1 spectral range. Not all absorbance wavenumbers are relevant to the analysis of biological material. Given the interdependence of these wavenumbers and the selective presence of relevant ones in biological material, techniques like PCA and 2D-COS are useful for data exploration. The complexity of biological tissue FT-IR data makes the incorporation of statistical methods beneficial for improved understanding of the FT-IR data (Marcott et al., 2014; Ali et al., 2018). Through the application of chemometric data analysis, it is possible to discern correlations among functional groups and visualize the distribution of multiple functional groups across the granule. This approach enhances the ability to identify and understand the relationships among different chemical constituents within the studied samples.

The utility of techniques such as PCA and 2D-COS extends beyond only FT-IR microscopy, as it can be applied for any spectroscopic measurement method such as Raman microscopy, Imaging Mass Spectrometry and Magnetic Resonance Imaging, among others (Gowen et al., 2015). Their versatility lies in their capacity to unravel complex datasets, identify correlations, and provide insightful visualizations not confined to a single analytical domain. This underscores their broader importance as powerful tools for comprehensive data exploration and interpretation across various scientific disciplines.

In the context of similar biological material compositions, differentiation between regions inside mapped areas is intricate, and complicated by the convolution of absorbance bands from overlapping functional groups. The penetration depth (up to 2 µm), as well as the spatial resolution of 1.56 µm in this FT-IR micro-spectroscope poses challenges in attributing specific observations to individual microorganisms, making e.g. microbial identification less straightforward compared to, for instance, laser microscopy (Gowen et al., 2015). It is also due to these limitations, that distinguishing between intracellular and extracellular components becomes challenging. Validation of observed polymers as part of the extracellular matrix, rather than the intracellular space, requires complementary techniques. In the future, it would be beneficial to perform validation studies using molecular imaging techniques of microbial composition i.e. FISH or with chemical or biological (lectin) staining methods. Nonetheless, this non-destructive imaging method could be a powerful imaging method for imaging the EPS distribution across the granular sludge.

#### Conclusion

In this study, FT-IR micro-spectroscopy was used to explore and visualize the EPS spatial distribution inside anaerobic granular sludge. FT-IR micro-spectroscopy is an untargeted imaging method that can measure a broad range of polymers simultaneously. With the single-point absorbance it was possible to create an absorbance height distribution for multiple different functional groups corresponding to proteins, polysaccharides, lipids, sialic acid and phosphorylated/sulfated polymers. Due to the complexity of the extracellular matrix, chemometric techniques were implemented to help elucidate the EPS composition. Principal component analysis made it possible to visualize multiple functional groups at the same time, which helped in distinguishing between regions that had a high absorbance height in similar functional groups. Two-dimensional correlation spectroscopy allowed for the identification of correlated functional groups. In this study we show that FT-IR micro-spectroscopy is a powerful complementary technique for EPS analysis and visualization. The combination of this non-destructive imaging method with other molecular imaging techniques would provide valuable insights in the role of specific polymers inside the extracellular matrix in anaerobic granular sludge.

## Acknowledgements

This work was financially supported by the SIAM Gravitation Grant 024.002.002, The Netherlands Organization for Scientific Research.

## Author contributions

SB and YL conceived and designed research. SB and CH conducted experiments and analysed the data. SB wrote the original manuscript with contributions/revisions from ML, YL and DS. All authors read and approved the manuscript.

## Preprint

The manuscript is deposited as pre-print in *bioRxiv*.

