## Supplemental information S1 for "FT-IR micro-spectroscopy for imaging the extracellular matrix composition in biofilms"

**MATLAB script for FT-IR imaging**

```
clearvars

% Load fsm data obtained from PerkinElmer frontier 400
micro spectroscope
% Data consists of an X Y Z block where X and Y are
location parameters and Z is the absorbance spectrum on
each X,Y location.
% Unfold the Z data block from [X,Y] to [X.Y,1] to ease
processing.
[data, xAxis, yAxis, zAxis, ~] =
fsmload('Path/To/Filename/Filename.fsm');
spectrum =
zeros(length(xAxis)*length(yAxis),length(zAxis));
n = 1;
for i = 1:length(xAxis)
    for j = 1:length(yAxis)
        spectrum(n,:) = data(i,j,:);
        n = n + 1;
    end
end

% Change transmittance to absorbance
abs_sp = 2 - log10(spectrum);
clear data spectrum

% Create a subset for region of interest (3500 to 800 cm-1)
sp = abs_sp(:,250:end-55);
zAxis_n = zAxis(250:end-55);
clear zAxis abs_sp n

% Smoothing (Savitsky Golay)
sp = smoothdata(sp(:,:),'sgolay',10);

% Simple baseline correction based on first order to
correct baseline drift
a = (sp(:,1) - sp(:,end)) ./ (3500 - 800);
corr = a .* zAxis_n;
bl_sp = sp(:,:) - corr(:,:);
bl_sp = bl_sp - bl_sp(:,1);
bl_sp = abs(bl_sp);
bl_sp = normalize(bl_sp,1,'range',[0 1]);
clear a corr s

% Filter out bad data points
a = mean(bl_sp,2);
BL_SP = a > 0.3;
bl_sp = bl_sp .* BL_SP;
```

```
% Plot average absorbance data
b = reshape(a,length(yAxis),length(xAxis));
b = b';
figure
image(xAxis,yAxis,b,'CDataMapping','scaled');
truesize
clear a

% Principal component analysis
[coeff,score,latent,tsquared,mu,explained] = pca(bl_sp);
% Plot the coefficient spectra/loading spectra of the first
5 PCs
figure
plot(zAxis_n,coeff(:,1),'r',zAxis_n,coeff(:,2),'g',zAxis_n,
coeff(:,3),'b',zAxis_n,coeff(:,4),'k',zAxis_n,coeff(:,5))
set(gca,'XDir','reverse');
legend('PC1','PC2','PC3','PC4','PC5');

% Imaging absorbance of single absorbance peaks
p1 = 1630; p2 = 1030; p3 = 2920; p4 = 1726; p5 = 1206;
C2 = zeros(192,192,5);
abs_p = zeros(192,192);
peaks = [ p1; p2; p3 ;p4;p5];
for p = 1:5
    for m = 1:length(xAxis)
        abs_p(m,:) =  bl_sp((m-
1)*length(yAxis)+1:m*length(yAxis),find(zAxis_n==peaks(p)))
;
    end
    C2(:,:,p) = abs_p;
end

% Imaging three functional groups at the same time
C3 = normalize(C2,1,'range');
figure;
image(xAxis1,yAxis1,data1);
hold on
image(xAxis,yAxis,C3(:,:,[1 4 5]));
title('polymers');
hold off

% Imaging PCA
pc_score = zeros(length(xAxis),length(yAxis));
part1 = score(1:length(xAxis)*length(yAxis),:);
C = zeros(length(xAxis),length(yAxis),3);
Cn = zeros(length(xAxis),length(yAxis),3);
color = [1,2,3]; % PC's chosen
for p = 1:3
    for m = 1:length(xAxis)
```

```
        pc_score(m,:) = part1((m-
1)*length(yAxis)+1:m*length(yAxis),color(p));
    end
    C(:,:,p) = pc_score;
end
C(C<0)=0;
% Normalize
Cn = normalize(C,1,'range');
b(b<0.3)=0; b(b>0.3)=1;
% Create image
figure;
image(xAxis1,yAxis1,data1);
hold on
image(xAxis,yAxis,Cn.*b);
hold off

% Two-D COS
A_av = mean(bl_sp,1);
A_vk = bl_sp - A_av;
A_vk(:,330:850)=0;
dot = cov(A_vk);
diago = diag(dot);
figure;
plot(zAxis_n,diago)
figure;
contourf(zAxis_n,zAxis_n,dot,'EdgeColor','none');
```
