## Supplemental figures S1-S4 for "FT-IR micro-spectroscopy for imaging the extracellular matrix composition in biofilms"

**Supplemental information**

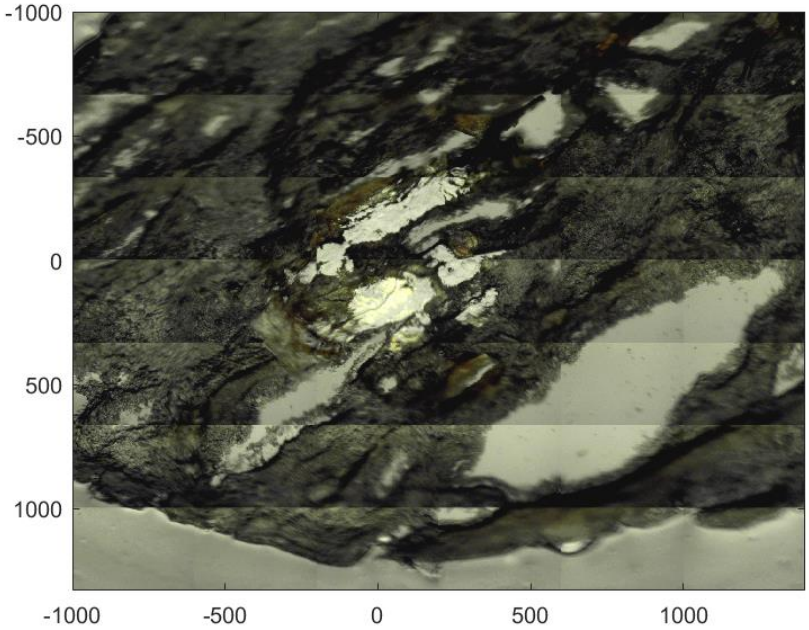


Figure S1. Sliced anaerobic granule used for FT-IR micro-spectroscopy. X and Y axis display distance from origin in μm.

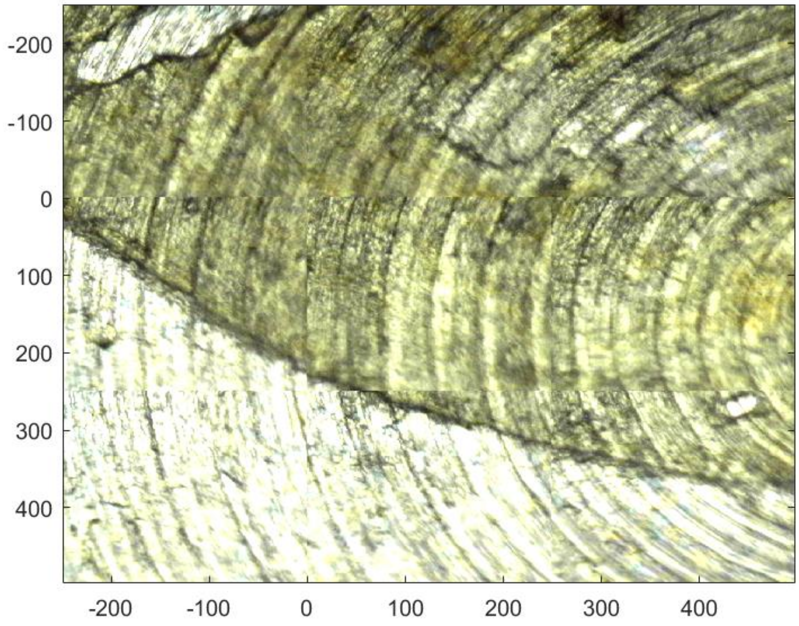


Figure S2. Sliced anaerobic granule used for FT-IR micro-spectroscopy. The background seen here is a metal wafer, hence the rings. X and Y axis display distance from origin in μm.

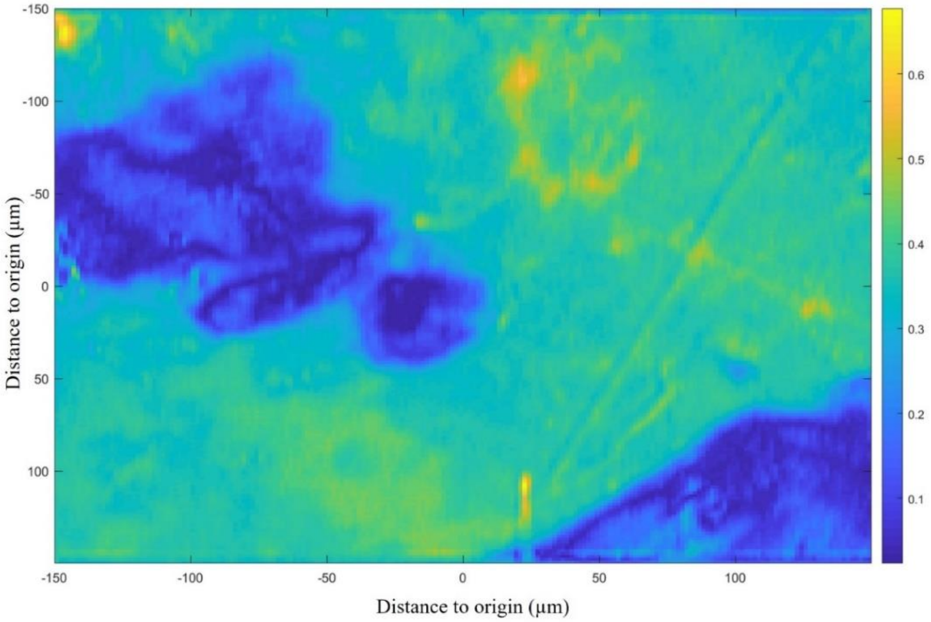


Figure S3. Average absorbance height (a.u.) of the FT-IR spectrum measured for granule (S1) for a 300 by 300 μm area.

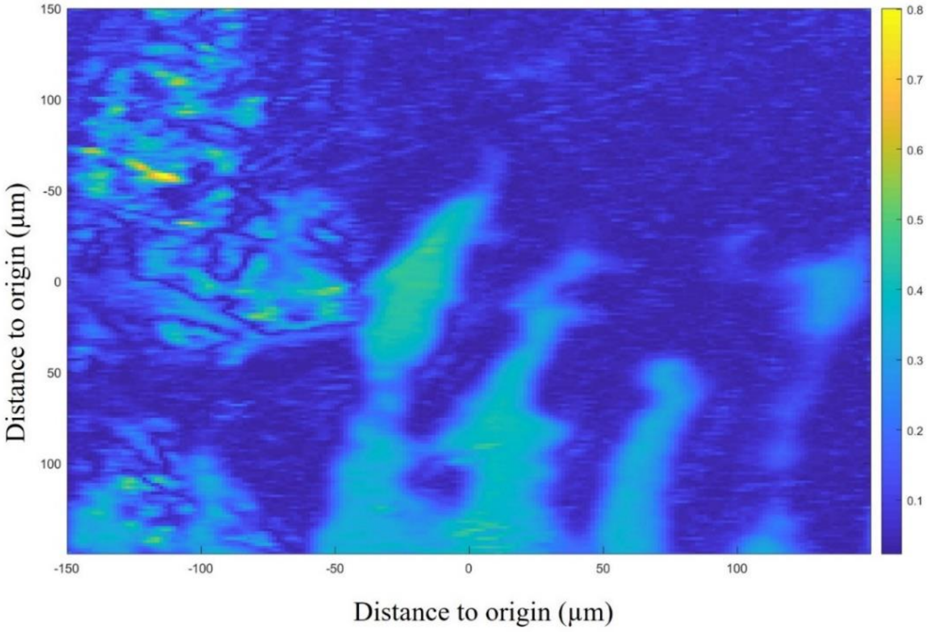


Figure S4. Average absorbance height (a.u.) of the FT-IR spectrum measured for granule (S2) for a 300 by 300 µm area.
